# The genomic geography and evolution of clusters of tandemly duplicated genes in the human and mammal genomes

**DOI:** 10.1101/504498

**Authors:** Juan F. Ortiz, Antonis Rokas

## Abstract

Clusters of duplicated genes (CTDGs) are nearly ubiquitous in life’s genomes, and are associated with several well-known gene families, such as olfactory receptors, zinc fingers, and immunity-related genes, as well as with several highly variable traits, including olfaction, body plan architecture, and pathogen resistance. However, these observations are usually anecdotal, restricted to specific cases, and lacking evolutionary context. In this study, we use a robust statistical approach to characterize the CTDG repertoire and analyze the distribution of CTDGs across 18 mammal genomes, including human. We found that, on average, 18% of the genes in each species are parts of CTDGs. Although genes in CTDGs are enriched for several biological processes, these tend to be involved in the interactions between the organism and its environment. We further found that mammalian CTDGs are not uniformly distributed across chromosomes and that orthologs of the human chromosome 19 are among the most clustered chromosomes in nearly all mammalian genomes analyzed. We also found evidence that the human chromosome 19 was formed by a fusion event that occurred before the diversification of the rodent and primate lineages and maintained its high density of CTDGs during its subsequent evolution. Finally, using chromosome-level alignments across mammalian genomes, we show how the syntenic regions of the human chromosome 19 have been shrinking, increasing their gene density and possibly increasing the compactness of its CTDGs. These results suggest that CTDGs are a major feature of mammalian genomes and provide novel insights into the origin and evolution of regions with unusually high densities of CTDGs.

## Introduction

Clusters of duplicated genes (CTDGs) are a familiar feature of genomes across the tree of life. CTDGs have been associated with phenotypic diversity in a wide variety of traits and organisms, such as in cellulose degradation in bacteria (Liebl et al., 1996), copper resistance in yeasts (Karin, et al., 1984), antimicrobial protection and venom in mammals and reptiles (Whittington et al., 2008), and pathogen resistance in plants (Michelmore, Meyers, 1998). The classic example of how CTDGs are involved in the generation of phenotypic diversity are the Hox gene clusters found in diverse metazoans; variation in the function and regulation of the Hox genes is one of the major contributors to the diversity of metazoan body plans (Garcia-Fernández, 2005; Lemons, 2006). Similarly, the gene duplications, conversions, and functional divergence of genes in the globin CTDGs in vertebrates are responsible for the functional diversity in the binding and transport of oxygen exhibited by members of this gene family (Hardies, Edgell, Hutchison, 1984; Runck, Moriyama, Storz, 2009).

One of the major dimensions of variation in CTDGs across genomes is the number of genes contained in a cluster. For example, the CTDGs formed by olfactory receptors (ORs) can be highly variable, from a single OR and the absence of a CTDG in elephant sharks, to thousands of ORs and 345 CTDGs in mouse (Niimura, 2009; Niimura et al., 2014). A similar trend is exhibited by genes in Killer Cell lectin-like receptor (KLRA) CTDGs; whereas only one KLRA pseudogene is found in human chromosome 12, 11 and 25 functional KLRA genes (plus 9 and 4 pseudogenized copies) are found in the syntenic regions of mouse chromosome 6 and rat chromosome 4, respectively (Kelley, Walter, Trowsdale, 2005).

In the human genome, in addition to the above mentioned OR, Hox, and globin examples, CTDGs are also known for gene families whose functions have been implicated in diverse processes, such as brain development (e.g., Protocadherin; Noonan, et al., 2004), immunity (e. g., KIR and Siglec; Martin et al., 2000; Cao et al., 2009), and pregnancy (e.g., Psg; Rawn, Cross, 2008). Interestingly, some of these CTDGs are known to be near each other in the genome. By far the most conspicuous aggregation of CTDGs to a single genomic region is human chromosome 19, which is characterized by an unusually high density of CTDGs (Grimwood et al., 2004). For example, zinc finger genes (Lukic, Nicolas, Levine, 2014), cytochrome P450 genes (Hoffman et al., 1995), Siglec genes (Cao et al., 2009), and gonadotropin beta-subunit genes (Nagirnaja, 2010) all exist as CTDGs in chromosome 19.

Although many human CTDGs have been well-characterized (Lukic, Nicolas, Levine, 2014, Than et al., 2014, Teglund et al., 1994, Hoffman et al., 1995; Cao, Crocker, 2011; Nagirnaja et al., 2010; Lindahl et al., 2013) and the high density of CTDGs in chromosome 19 is well-established (Grimwood et al., 2004), a systematic study of the distribution of CTDGs in humans and the degree to which they are conserved in other mammals is lacking. For example, are CTDGs a common feature of the human genome, and more generally of mammalian genomes, or a rare one? Are CTDGs randomly distributed across human and mammalian chromosomes? How conserved are human CTDGs and CTDG neighborhoods in other primates and mammals?

To address these questions, we comprehensively characterized the genomic geography and evolution of CTDGs in the human and select high-quality mammalian genomes. We found that CTDGs are common genomic features, being present in all mammalian genomes examined and significantly associated with biological processes, such as olfaction, signaling, and immunity. Furthermore, mammalian CTDGs are not uniformly distributed; the most spectacular case of this deviation is the human chromosome 19 and its orthologs in mammals, which are 1.75 times more densely populated with CTDGs than the average mammalian chromosome. Interestingly, we found that human chromosome 19 formed from the fusion of two ancestral chromosomes, only one of which was CTDG-rich. By exploring the relationship between gene family dynamics and CTDG membership, we observed that gene number expansion and contraction events are concentrated at the base of the mammalian phylogeny, and in gene families with low (0 – 25%) to medium low (25% - 50%) fractions of their gene members being parts of CTDGs, suggesting that most CTDGs present in mammalian genomes stabilized their gene numbers prior to the diversification of the major mammalian lineages.

Aside from mapping the genomic geography of CTDGs across mammals and tracing their evolution, this work provides a methodological framework for the genome-wide evolutionary study of CTDGs as well as support for the idea that CTDGs are biologically relevant features of genomes. Our data show that CTDGs are not mere statistical artifacts distributed at random; rather, they are concentrated in some regions, with human chromosome 19 being an extreme such example. Furthermore, CTDGs can be both diverse in gene numbers and density at the interchromosomal level, but also stable across (chromosome) orthologs. These observations open interesting avenues of research. For example, understanding how evolutionary processes modulate the diversification and stabilization of CTDGs could yield insights into how the evolution of genes, gene families, and chromosomes are related.

## Methods

### Data acquisition

The genomes of human, select primates (*Callithrix jacchus*, marmoset; *Chlorocebus sabaeus*, green monkey; *Gorilla gorilla*, gorilla; *Homo sapiens*, human; *Macaca mulatta*, macaque; *Nomascus leucogenys*, gibbon; *Pan troglodytes*, chimpanzee; *Papio anubis*, baboon; *Pongo abelii*, orangutan), and select mammals (*Bos taurus*, cow; *Canis familiaris*, dog; *Equus caballus*, horse; *Felis catus*, cat; *Monodelphis domestica*, opossum; *Mus musculus*, mouse; *Ovis aries*, sheep; *Rattus norvegicus*, rat; *Sus scrofa*, pig) were downloaded from the REST Ensembl service (https://rest.ensembl.org, https://rest.ensemblgenomes.org; date of access: Aug 10, 2018). For each species, gene annotations, proteome sequences, and chromosome lengths were downloaded from Ensembl, keeping only genome contigs with more than 10 genes and longer than 10 Kbp (see table S1 for genome assembly version information).

### CTDG *identification*

To identify clusters of tandemly duplicated genes, we used a modified version of the CTDGFinder software (Ortiz, Rokas, 2017). Instead of using blast results from a given sequence query, CTDGFinder was modified to use the Panther database of protein families (Mi et al., 2017). Briefly, all the HMM profiles of protein domains in the Panther (v. 12; Mi et al., 2017) database were first used to annotate the proteomes of each genome using the HMMer sequence search software, version 3.1b2 (Eddy, 2011). HMMer subject-query hits with less than 30% mutual coverage were discarded.

Next, for each protein domain identified in a genome, the chromosomal coordinates of all the proteins in each chromosome (or genomic scaffold) that contained that protein domain were processed by the MeanShift algorithm (Comaniciu, 2002) to obtain CTDG candidates (i.e., the genomic region(s) with the highest densities of that domain). Statistical significance, i.e., whether a CTDG candidate was considered a CTDG, was evaluated by comparing the number of proteins belonging to a Panther family in the genomic region identified by the MeanShift algorithm against an empirical distribution obtained by counting the highest number of genes that belong to a Panther family in each of 1,000 randomly sampled genomic regions from the same genome of length equal to that of the CTDG candidate. A CTDG candidate was considered a CTDG if its number of genes containing the belonging to a Panther family was equal of greater than the 95^th^ percentile value of the distribution generated by the examination of 1,000 randomly sampled genomic regions. In that way, CTDGFinder analysis resulted in clusters containing proteins that shared at least one Panther family.

The next step consisted of collapsing CTDGs with overlapping coordinates, so that potential nested CTDGs were collapsed into a single CTDG. As a result, each CTDG may contain genes that belong to one or more Panther families. Finally, we rerun the MeanShift algorithm to further refine CTDG boundaries.

As CTDGFinder uses a statistical approach to identify clusters of tandemly duplicated genes, some CTDGs also contain genes that do not share homology with the set of tandemly duplicated genes that defines the cluster (e.g., a 5-gene CTDG may be comprised of 4 duplicated genes and 1 unrelated gene). Those unrelated genes were not considered when calculating various gene-based metrics (e.g., total gene length, number of genes), but were considered for the calculation of various cluster-based metrics (e.g., length of clustered regions, percentage of chromosome length spanned by CTDGs), as they contributed to the length of the entire cluster.

### GO enrichment

To explore any potential biases in the gene functional categories present in CTDGs, we performed GO enrichment analyses. Per gene GO annotation from each species was downloaded from the BioMart service on Ensembl (last accessed Aug 10, 2018), and the gene ontology files were downloaded from geneontology.org (last accessed April 29, 2017). GO enrichment analysis was performed by comparing the distributions of clustered genes in each chromosome against all protein-coding genes in the genome for human and mouse using the python module Goatools (https://doi.org/10.5281/zenodo.31628). Statistical significance was assessed using the Benjamini/Hochberg false discovery rate method (Benjamini, Hochberg, 1995) with a significance cutoff of 0.005.

### Chromosome orthology calling

To compare the chromosomal distribution of CTDGs across species, we inferred orthologous protein-coding regions for all human chromosomes. To do so, we first downloaded Ensembl’s orthology assignments for all the genes in the genomes we examined using Ensembl’s REST service (www.rest.ensembl.org; last accessed March 10, 2018). Next, given two chromosomes (A, B), one from the human genome and the other from a different mammalian genome, we calculated the proportion of genes in chromosome A that have orthologs in chromosome B (O_A,B_). Similarly, we calculated the proportion of genes in chromosome B that have orthologs in chromosome A (O_B,A_). Finally, if at least one of O_A,B_ or O_B,A_ was greater than 0.25, we considered chromosomes A and B to be orthologs.

### Clusteredness estimation

The distribution of CTDGs is not uniform across chromosomes. To compare the distribution of CTDGs between species, we first quantified how clustered each chromosome is by calculating how different each chromosome’s distribution of CTDGs was to all the other chromosomes in a given genome. Examination of the ranking of each chromosome in a given genome based on its proportion of CTDGs provided an assessment of how clustered each chromosome was in that genome. We call this ranking the clusteredness of a chromosome.

The clustering pattern of a chromosome was measured by calculating the distribution of clustered genes in non-overlapping 10Kb windows on each chromosome. That distribution was normalized, so that it could be treated as a probability mass function (PMF). To compare the clustering pattern of two chromosomes, we calculated the Jensen-Shannon distance (JSD) (Lin, 1991) from their PMFs. The all versus all pairwise JSDs were then used to generate a graph, where each chromosome corresponds to each node in the graph, and each JSD between each pair of chromosomes corresponds to each edge in the graph. We then used the farness centrality measure (FCM) (Marsden, 2002) to measure how distant a chromosome was to the rest of the chromosomes in a genome. Briefly, the FCM for a given chromosome A was computed by adding all the JSDs connecting to A, and then normalized by dividing the raw FCM value by the number of nodes minus one. The clusteredness rank of a chromosome was computed by sorting the chromosomes according to their normalized FCM values, so that the chromosome with rank 1 was the chromosome with the most different clustering pattern relative to the rest of the chromosomes.

### Chromosome length evolution

The observation that certain chromosomes remained highly clustered through evolutionary time proved to be an interesting aspect of our analysis. To gain further insight into the evolutionary processes underlying this conservation, we examined the dynamics of CTDG expansions and contractions of all human chromosomes specifically along the path from the mammalian common ancestor to the human lineage tip in each chromosome. Multiple chromosome alignments for all human chromosomes were downloaded from Ensembl, using their “25 mammals” EPO alignments (Vilella et al., 2008), including inferred syntenic sequences for internal nodes in the chromosome alignment tree accompanying each alignment (alignment tree). The dynamics of contractions and expansions during the mammalian history of each human chromosome were examined by performing linear correlations between the length of the alignment at each node on all alignment trees (for each human chromosome), and the relative age of each node (defined as the distance from the root of the tree, to the most recent common ancestor of the focal node and the human tip, normalized by the length of the tree).

### Gene copy dynamics

It has been well established that gene duplication and loss are the main processes by which gene families evolve (Lynch, Force, 2001). Since CTDGs are typically the evolutionary end result of a series of gene duplication and loss events, we investigated the gene number dynamics of clustered genes. The OMA standalone version (Altenhoff et al., 2015) was used to identify hierarchical orthology groups (Altenhoff et al., 2013) of the clustered genes found in this study. Gene gain and loss events were estimated using the PyHam python module, using the mammalian species phylogeny obtained from the TimeTree database (Kumar et al., 2017).

## Results

### CTDGs are abundant and non-uniformly distributed across mammalian genomes

To get a comprehensive picture of the distribution of clustered genes across species, we used CTDGFinder to identify CTDGs in select high-quality mammal genomes (see methods). On average, 16.0% (standard deviation 4.3%) of the genes in a mammalian genome are clustered. The genome with the smallest fraction of clustered genes was that of pig (6.2%) and that with the largest was mouse (24.3%). Examination of the distribution of CTDGs in the context of the mammalian phylogeny reveals a small decrease in the fraction of clustered genes in the primate lineage (Fig. 1A; average percentage of genes clustered in primates = 14.4%, average in non-primate mammals = 17.7%). Our GO enrichment analysis shows that the biological processes most commonly enriched in clustered genes in human and mouse are related to olfaction, signaling, and immunity (Fig. S1; Table S3).

**Figure 1.**
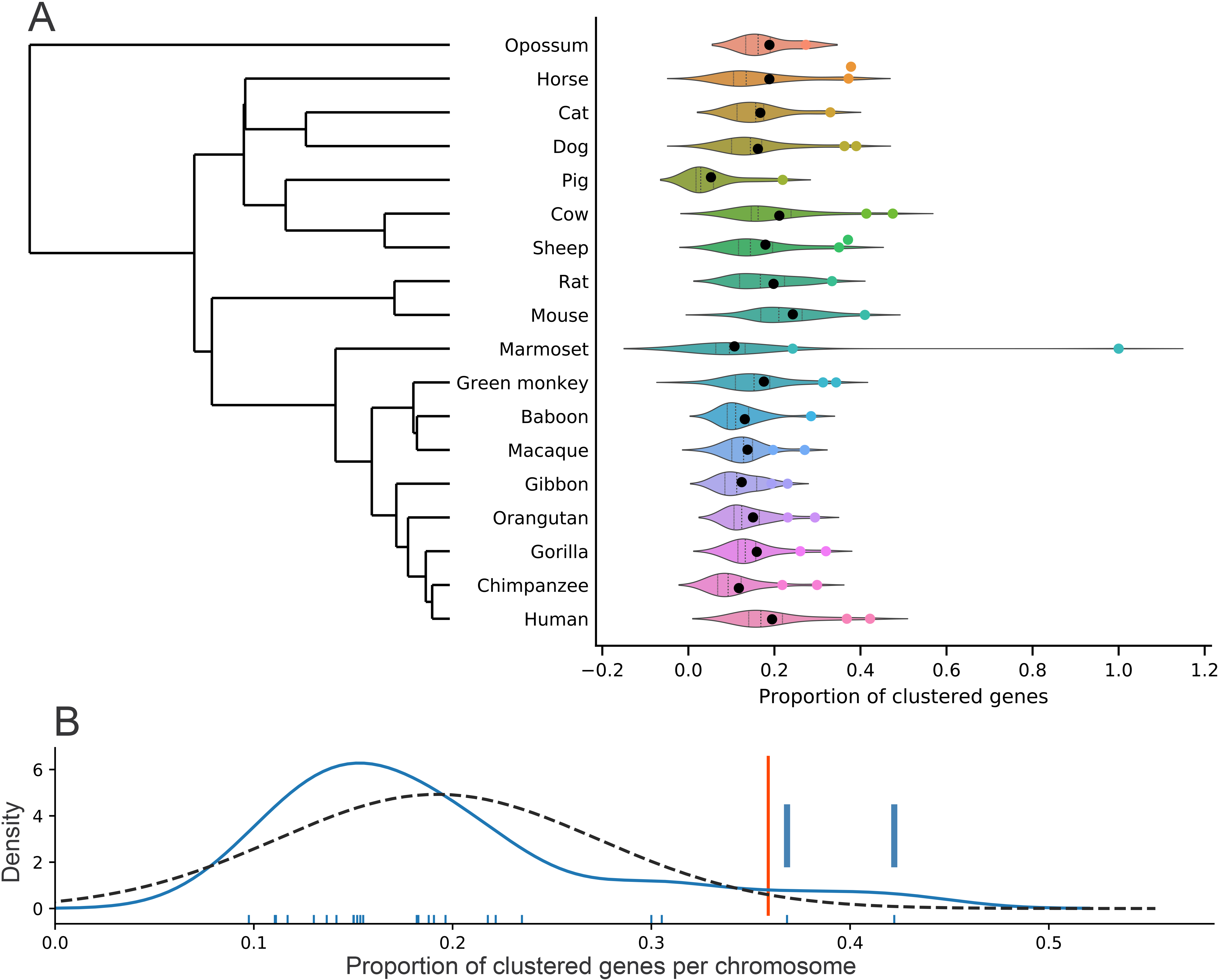
A large proportion of genes in mammalian genomes belongs to clusters of tandemly duplicated genes (CTDGs). (A) Distribution of CTDGs across the mammalian phylogeny. Violin plot: distribution of the proportion of clustered genes across chromosomes. Lines inside the plots show the interquartile ranges. Black: Mean proportion of clustered genes in each species. Colored dots: chromosomes with higher proportion of clustered genes than the 95^th^ percentile. (B) Detail of the distribution of the proportion of clustered genes in the human genome. Ticks: data points for each chromosome. Blue curve: KDE estimate. Dashed curve: Fitted normal distribution. Red line: 95^th^ percentile. Blue bars: Chromosome with higher proportion of clustered genes than the 95^th^ percentile.

Importantly, CTDGs are not uniformly distributed across chromosomes (Fig. 1A, S2). In all species, there are at most three chromosomes that seem to have a higher fraction of clustered genes compared with the rest of the chromosomes in each genome (Fig. S2). This observation is of particular interest, because it suggests that extreme clustering patterns could have been conserved in orthologous chromosomes during the evolution of mammals, giving rise to the idea that the genomic organization of CTDGs is subject to selection, and serves as a relevant hypothesis to be tested in this study.

### One fifth of human genes are part of CTDGs

To infer the distribution of clustered genes in the human genome, we used the CTDGFinder algorithm to examine the frequency of CTDG occurrence across all human chromosomes. We found that 3,821 / 19,665 (19.4%) human genes belong to 961 CTDGs, which collectively occupy 163.5 Mbp (5.3%) of the human genome. The average length of genes belonging to CTDGs is 18 Kbp, which is significantly lower than an average length of 53 Kbp for non-CTDG genes (Mann-Whitney P-value: 6.6×10^−256^). Consistent with our previous results (Ortiz & Rokas, 2017), the CTDGFinder software accurately identified several well-known CTDGs (Fig. 2). For example, the Siglec (Cao et al., 2011), Hox (Krumlauf, 1994), and globin (Efstratiadis et al., 1980) CTDGs were all correctly identified in the human genome. Similarly, the HLA genes (Tao et al., 2015) were found in 3 CTDGs composed of 12, 4, and 19 genes in chromosome 6; the protocadherin CTDG was found in chromosome 5 with 55 genes (Noonan et. al., 2004); the previously reported ORs in chromosome 11 (Niimura, 2009) were recovered as 14 CTDGs that contained a total of 193 genes; and the histone cluster in chromosome 6 (Albig et al., 1997) was found in three CTDGs of 47, 4, and 15 gene duplicates. Interestingly, the longest CTDG in the human genome is the protocadherin CTDG (Wu, Maniatis, 1999) on chromosome 5 with 55 genes. The 4-gene beta-globin CTDG was found in embedded in an OR-rich CTDG on chromosome 11 (Bulger et al., 1999). The next largest CTDG, with 46 duplicates was a histone-containing CTDG on chromosome 6. A comprehensive table with the CTDGs (and their genes) for all mammalian genomes analyzed in this study, including the human genome, can be found in supplementary table S2.

**Figure 2.**
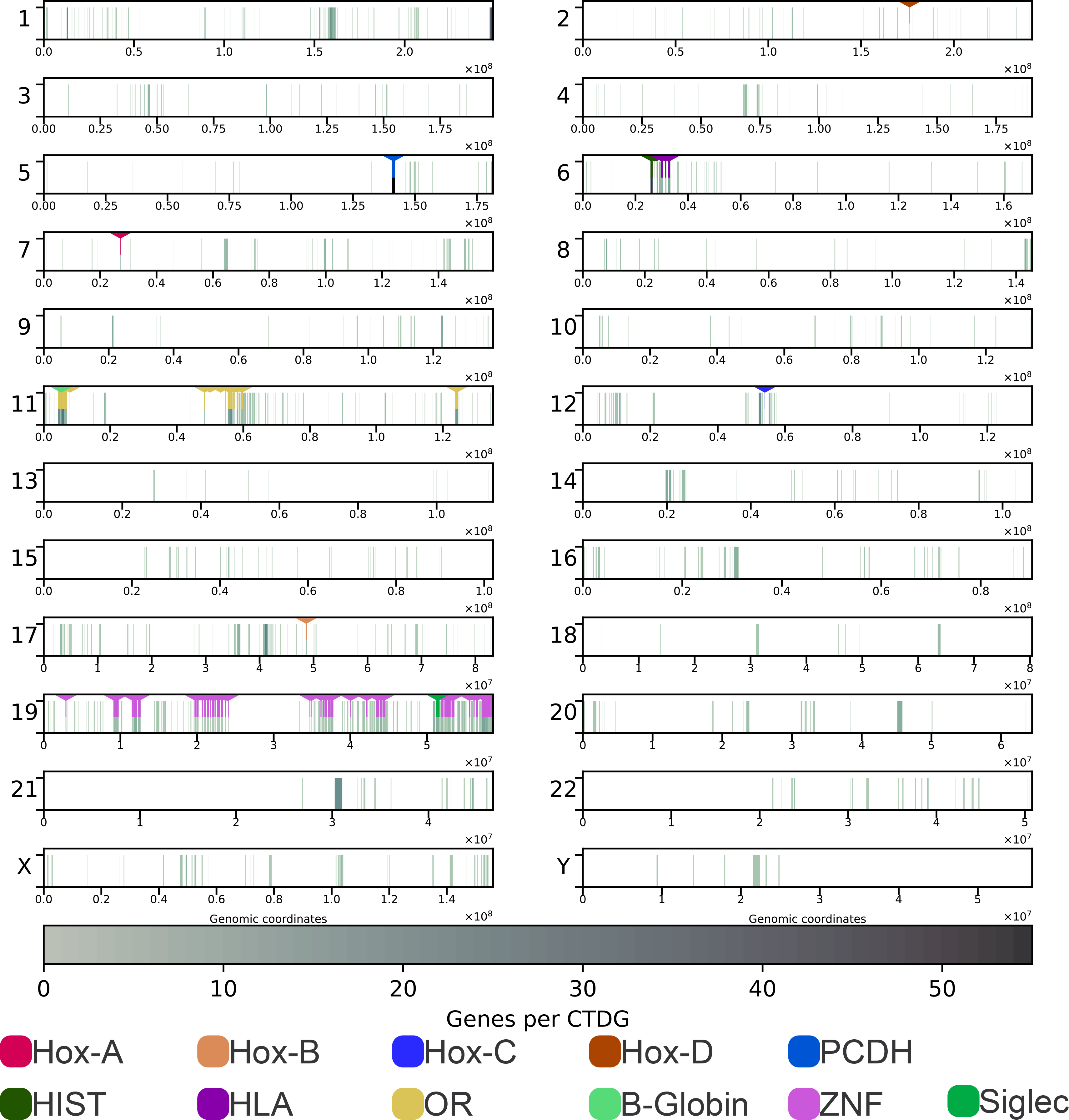
Distribution of CTDGs across all human chromosomes. Each plot represents a chromosome. The location and length of all CTDGs are shown as rectangles in the bottom section of each plot. Representative CTDGs are highlighted by a rectangle in the top section of the plots. An extended polygon is included in short CTDGs to improve visibility.

### Human chromosomes 19 and Y have the highest densities of CTDGs

To test if CTDGs are uniformly distributed across the human genome, we characterized each of the human chromosomes in terms of their number and total length of clustered genes, as well as the number and length of CTDGs. We found that on average, 5.92% of each chromosome’s length is covered by CTDGs and on average, 19.23% of genes in each chromosome belong to CTDGs. The two chromosomes that exhibit the highest percentages of clustered genes in the human genome are chromosome Y (19 clustered genes / 45 genes; 42.2%) and chromosome 19 (525 clustered genes / 1,431 genes; 36.7%) (Figs. 1B and 1C). The high percentage of clustered genes in chromosome Y is likely due to its small number of genes. Furthermore, the percentage of chromosome Y length spanned by CTDGs is 2.9%, which is below the average (5.9%) and on par with several other human chromosomes. The longest CTDGs in chromosome Y contain RNA-binding motif Y chromosome (RBMY) genes, which encode for nuclear proteins involved in spermatogenesis (Chai et al., 1998). Other CTDGs include deleted in azoospermia (DAZ) genes (Saxena et al., 1996), and testis-specific protein Y-linked (TSPY) genes (Vogel, Schmidtke, 1998).

In contrast to chromosome Y, whose high percentage of CTDGs appears to be at least in part a consequence of its small gene number, the much larger chromosome 19 appears to be genuinely densely populated with CTDGs. Specifically, not only more than one third of its genes belong to CTDGs (525 / 1,431), but also 32.0% of its length is composed by CTDGs, by far the highest of human chromosomes. For comparison, the chromosome with the second highest percentage of chromosome length spanned by CTDGs is chromosome 11, with only 12.1%. Furthermore, the percentage of the length of chromosome 19 that is spanned by CTDGs is higher than the 95th percentile of the data, and higher than two times the standard deviation of a normal distribution fitted to the percentage of chromosome length spanned by CTDGs in all the human chromosomes (Fig. 1B). These data suggest that CTDGs are not uniformly distributed across chromosomes, with chromosome 19 being by far the most notable deviation (Fig. 1B). The CTDG with most genes in chromosome 19 contains 19 zinc finger duplicated genes. In addition to these zinc finger CTDG, chromosome 19 also contains the KIR (Martin et al., 2000), kallikrein (Lindahl et al., 2013), Siglec (Cao et al., 2009), OR (Niimura, 2009), CEACAM (Cao et al., 2009), PSG (Teglund et al., 1994), and Galectin (Than et al., 2014) CTDGs.

### Gene family dynamics of CTDGs

To gain insight into when gene families associated with CTDGs experienced gene gain and loss events, we estimated gene duplication and loss events on genes that are parts of CTDGs. We found that most of the duplication events in the dataset preceded the diversification of eutherians. Interestingly, most gene gains and losses in clustered genes occurred in the rodent lineage (Fig. 3).

**Figure 3.**
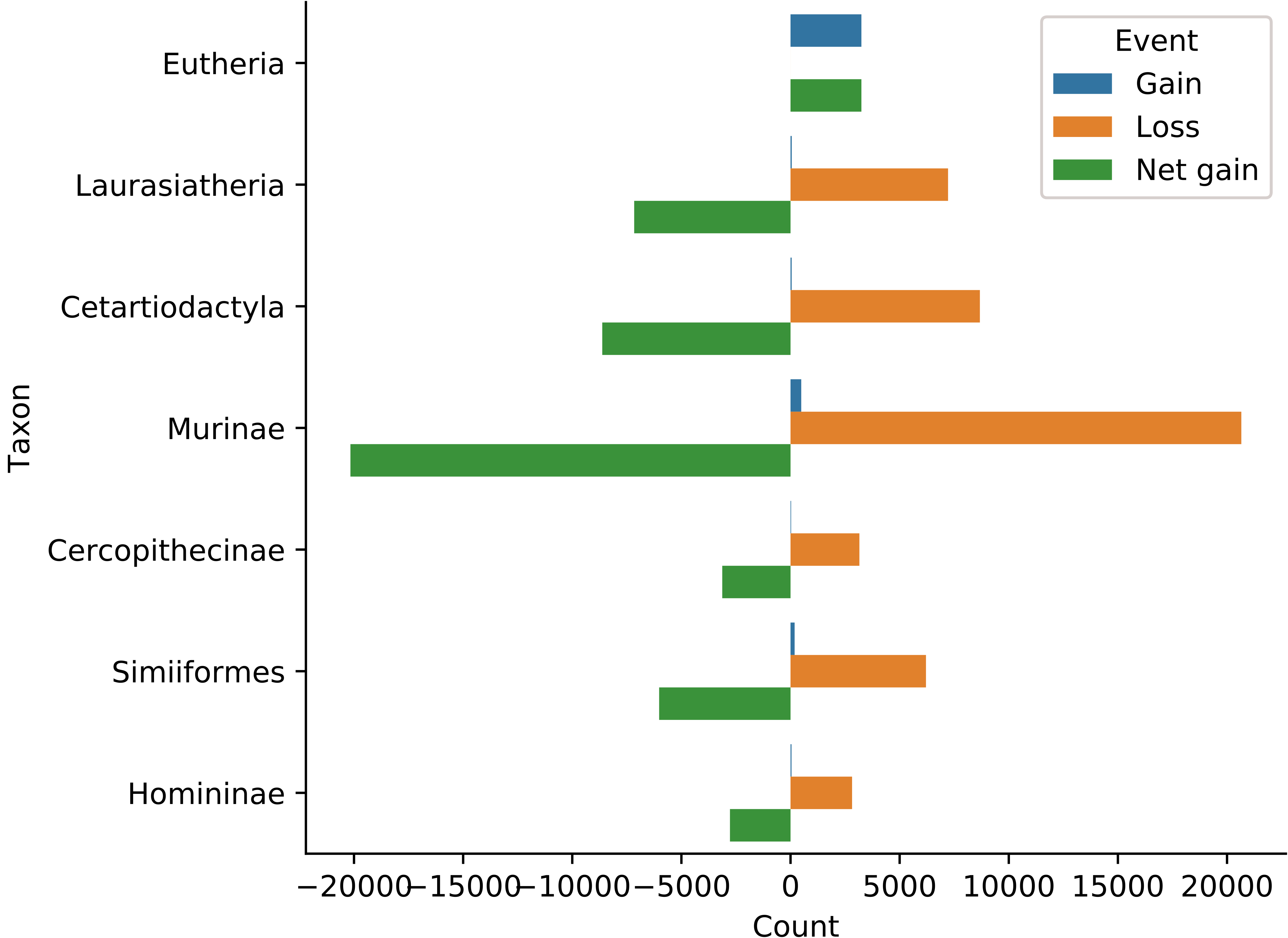
Gene family dynamics across the mammalian lineage. Each gene tree was given a cluster status depending on the proportion of clustered genes in them (low: less than 0.25, mid low: between 0.25 and 0.5, mid high: between 0.5 and 0.75, and high: more than 0.75). (A) Phylogeny of mammalian genomes used in this study. (B) Gene duplication events. (C) Gene loss events. (D) Net gene gains (gene duplications - gen losses).

### The extreme clustering of human chromosome 19 is highly conserved across mammals

Our finding that there are at most three chromosomes in each mammalian genome that contained a significantly higher fraction of clustered genes than the rest of the chromosomes raised the possibility that these highly clustered chromosomes are in fact orthologous and have maintained their clusteredness during the evolution of mammals. We found that in all the genomes studied here, except dog and pig, at least one of the five most clustered chromosomes is orthologous to the human chromosome 19 (hc19 ortholog). In the case of the dog genome, the two chromosomes with highest clusteredness rank are orthologs of the human chromosome 11, which is the third most clustered human chromosome (below chromosome 19 and chromosome Y) and is also known to be rich in CTDGs, including an OR CTDG (Niimura, 2009) and a fibroblast growth factor CTDG (Roy, Calaf, 2014). Most mammalian genomes contain a single hc19 ortholog that is highly clustered; that is the case for all primates, except gibbon, rodents, as well as horse (Fig. 4). Moreover, in genomes that contain two chromosomes considered by our cut-offs orthologous to hc19 (see methods), except gibbon, only one of these two chromosomes is highly clustered. The gibbon chromosomes are the exception, presumably because the gibbon genome has experienced a very large number of genomic rearrangements, especially around transposable elements (TEs), compared with other primates (Carbone et al., 2014). Given that CTDGs tend to be depleted in TEs (Grimwood et al., 2004), it is likely that CTDGs from the hc19 orthologs in gibbon were broken up by TE insertions, shuffled, or transposed to other chromosomes. In order to test if the highly rearranged genome affected the distribution of CTDGs in gibbon, we run phylogenetic independent contrasts (Felsenstein, 1985) on the mean fraction of chromosome length spanned by clusters, and the mean fraction of clustered genes across chromosomes in the primates in our dataset. We found that although gibbon has the lowest mean fraction of chromosome length spanned by CTDGs, it differs from marmoset (the second lowest value) in less than 0.002 phylogenetic independent contrast (PIC) units (mean PIC: 0.05; Fig 5.A). Similarly, the mean fraction of clustered genes per chromosome in gibbon shows little variation from the rest of the primates (Fig 5. B).

**Figure 4.**
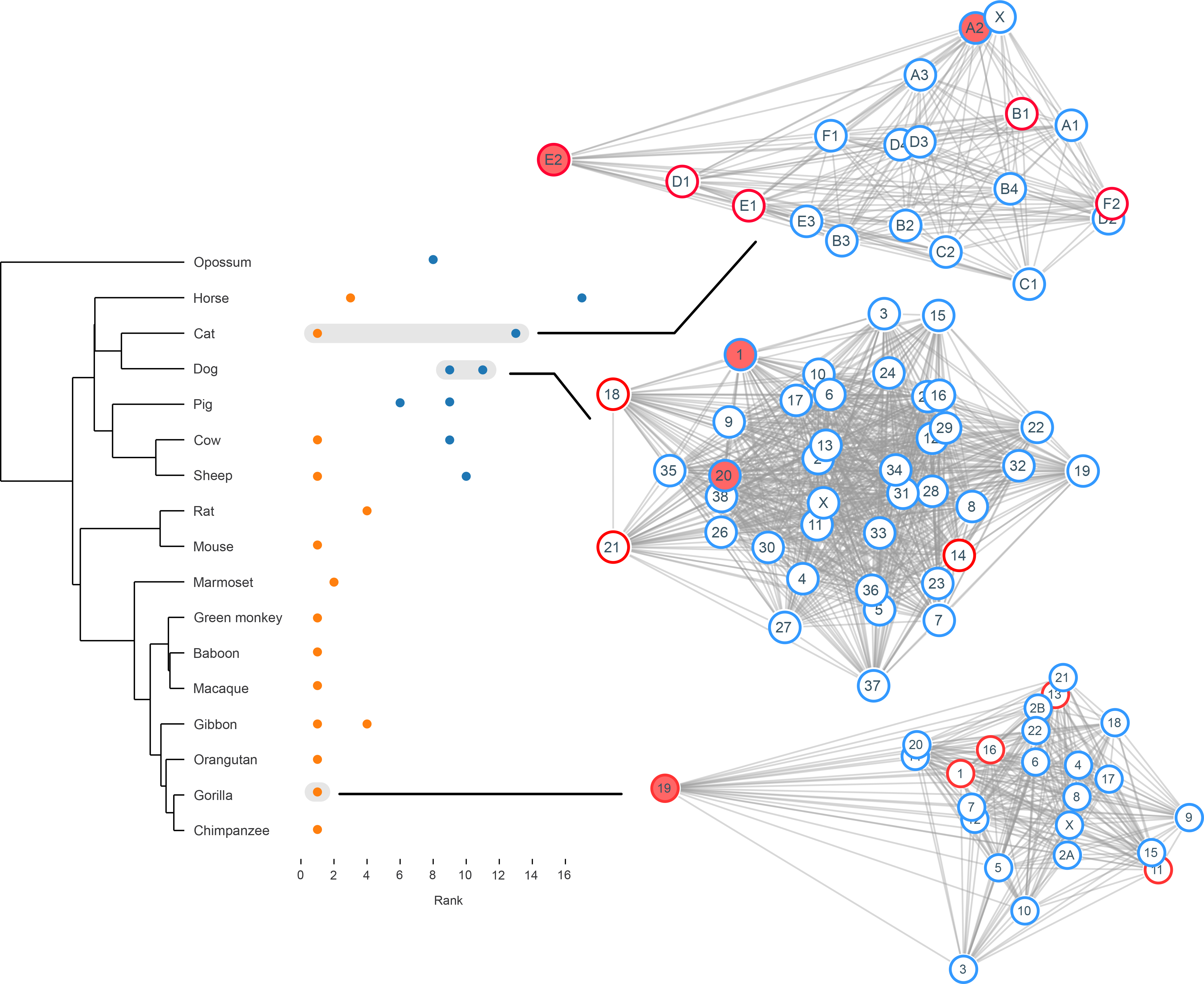
Clusteredness rank of the chromosome 19 orthologs on selected species. Orange: Chromosomes on the top 5 rank. Blue: Chromosomes on ranks above 5. To the right, network representations of selected species. Red border: Chromosomes in the top 5 rank. Pink: Human chromosome 19 orthologs. Edges are drawn using 1 - farness as weight.

**Figure 5.**
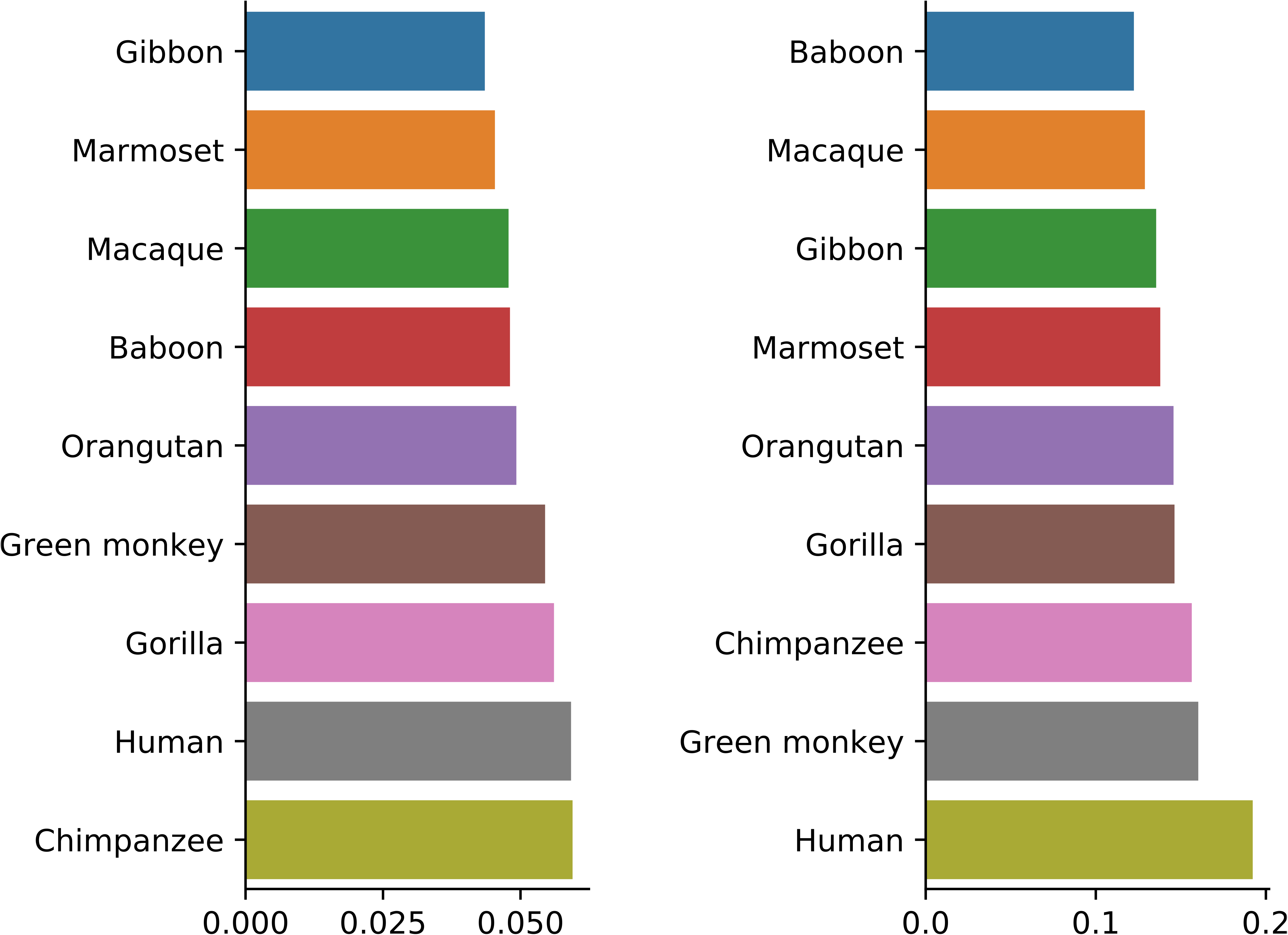
Phylogenetically corrected variation in CTDG length and composition across primates. A: Mean fraction of chromosome region spanned by CTDGs. B: Mean fraction of clustered genes per chromosome

To understand the evolution of clustering of human chromosomes, we used the EPO genomic chromosome alignments from Ensembl (Vilella et al., 2008), including inferred ancestral sequences, to estimate the relative time (in phylogenetic tree units) at which the syntenic regions of each of the human chromosomes across 26 mammals changed in size. By examining the change in the length of syntenic sequences (extant and ancestral) for each human chromosome in the context of the mammalian phylogeny, we inferred the rate at which each the orthologous regions of each chromosome shrunk or expanded (Fig. 6). Specifically, for all the human multiple chromosome alignments we built linear regressions of the aligned genomic ancestral sequences at each node and the ages of the nodes examined. The slopes of the regressions between changes in alignment length and evolutionary time are positive for all human chromosomes, which suggests that all human chromosomes, or more precisely all aligned regions of human chromosomes, have been shrinking (Fig. 6A). Furthermore, along with chromosome Y (Pearson R value = 0.17; Fig. 6D), chromosome 19 shows the largest slope (Pearson R value = 0.19; Fig. 6C), suggesting that human chromosome 19 has been shrinking at a higher rate than the rest of the human chromosomes. To test if the observed behavior was a byproduct of chromosome size, we carried out a correlation of the calculated slope against chromosome size. We found a strong correlation between slope and chromosome length, suggesting that chromosome length is a confounding variable in our calculations. However, both chromosome 19 an Y showed by far the highest slope of all human chromosomes, including when compared to other small human chromosomes, such as chromosomes 20, 21, and 22 (Fig. 6B).

**Figure 6.**
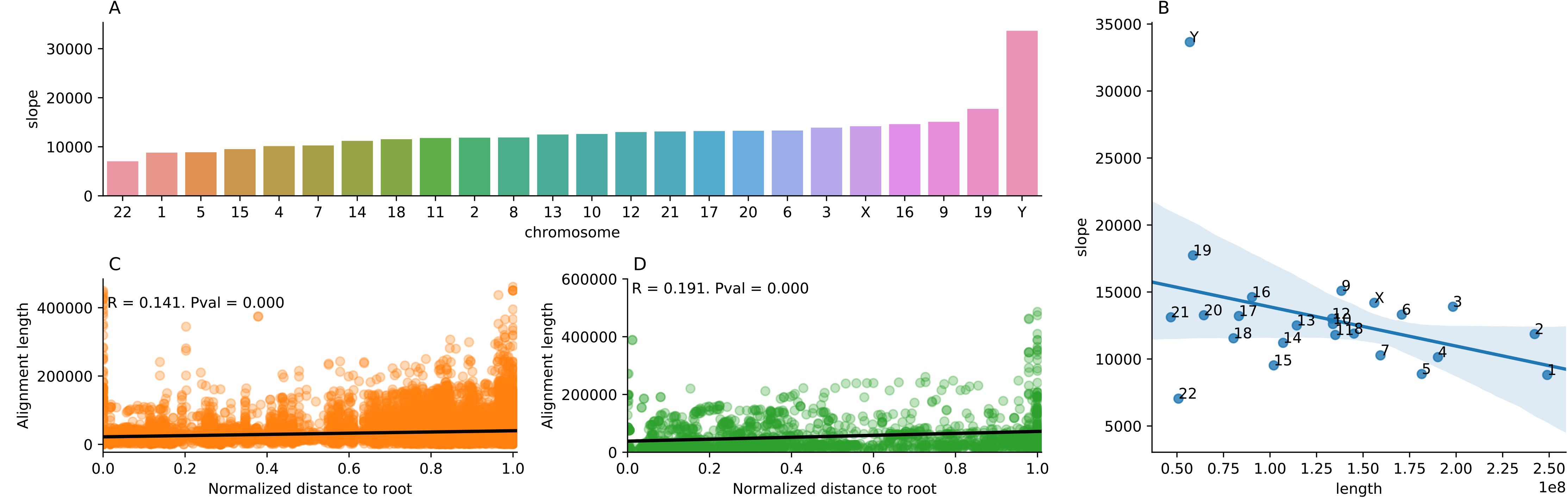
Rate of human chromosome shrinking in the mammalian lineage. A: slope calculated from correlations between the total alignment length of all EPO Ensembl alignments for each human chromosome and the normalized distance to the root of the tree of the EPO alignment at each node. B: Influence of chromosome length in the slope calculated in A. C: Example correlation for chromosome 19 (R: pearson R value; Pval: P-value). D: Example correlation for chromosome Y.

## Discussion

Historically, the discovery and description of CTDGs has been piecemeal and subjective (Duboule, 2007; Ortiz, Rokas, 2017). Although studies on subjectively defined CTDGs have been extremely useful and have greatly advanced our knowledge of their evolution and function (Alam et al., 2006; Lemons, 2006; Noonan, 2004; Than et al., 2014), a comprehensive investigation of CTDG evolution has so far been lacking. In this study, we provide the first systematic characterization of the genomic geography and evolution of CTDGs in mammalian genomes. Although it is hardly surprising that mammalian genomes harbor CTDGs, our findings suggest that CTDGs are highly prevalent – nearly one in five mammalian genes are part of CTDGs – and constitute common and conserved features of mammalian genomes.

CTDGs are not uniformly distributed across mammalian genomes; the same is true for the human genome (Fig. 2). Our results showcase that CTDGs can vary widely in terms of their gene density, length, and number of genes. For example, with a mean cluster length of 147 Kbp, the iconic Hox clusters in humans (Holland, Garcia-Fernández, 1996) (Fig. 2, chromosomes 2, 7, 12, and 17) are relatively small in size compared to a 1,402 Kbp long zinc finger CTDG in the same genome (Lukic et al., 2014) (Fig 2, chromosome 19). The human protocadherin CTDG (Noonan, 2004), which is 1,172 Kbp long, is very gene-dense, with 55 clustered genes; while a histone CTDG in chromosome 6 has a similar number of clustered genes (47), but it is only 783 Kbp. The diversity of CTDG length and gene density can be further appreciated in the 14 OR-containing CTDG in chromosome 11 on the human genome (Niimura, 2009) (Fig. 2, chromosome 11), which have sizes ranging from 30 Kbp to 917 Kbp. Moreover, while the longest CTDG (1.4 Mbp) has 10 gene duplicates, the second longest CTDG (1.2 Mbp) has 43 gene duplicates.

From the genomic landscape perspective presented here, we can visualize how chromosome 19 is especially CTDG-rich, with more than 13% of the clustered genes in the human genome being located there (Fig 2, chromosome 19). Furthermore, our study found that 36.7% of chromosome 19 genes are part of CTDGs, a substantial increase over the previously reported estimate of 30% (Dehal et al., 2001).

Importantly, CTDGs can contain more than one clusters; such is the case of the 5-gene globin cluster, which is located inside an 8-gene OR cluster (Bulger et al., 1999) (Fig 2. chromosome 11). In terms of the high physical linkage of their genes and their involvement in complex phenotypes, it could be said that CTDGs resemble supergenes (Schwander, Libbrecht, Keller, 2014). However, we did not see a high difference in recombination rate on sites inside and outside CTDGs (Fig. S3), with the expected exception of chromosome 6, where the MHC complex resides, which has been observed to be highly polymorphic, with high recombination observed inside the MHC cluster (Yeager, Hughes, 2006).

The high density of CTDGs in chromosome 19 puzzled us because the evolutionary maintenance of all these CTDGs “in place” likely required strong selection. By tracing the evolutionary history of chromosome 19 “clusteredness”, we found that chromosome 19 was formed by a chromosomal fusion before the diversification of primates and rodents (Fig. 3). Interestingly, we found that most of the CTDGs in the human chromosome 19 come from only one of the two proposed chromosome 19 orthologs, which points to a model where CTDGs have indeed been maintained in the mammalian lineage. Furthermore, examination of chromosome-level sequence alignments suggests that the syntenic regions of chromosome 19 have been shrinking (Fig. 6A). A caveat of our experiments is that we can only examine the evolution of the parts of chromosome 19 that are syntenic and alignable across species; thus, it is possible that the chromosome itself could be expanding if the amount of non-homologous sequence has increased during evolutionary time. That caveat notwithstanding, we hypothesize that the high density of CTDGs might have evolved by contractions in chromosome length.

Why would CTDGs in mammalian genomes appear to be so well conserved? One likely explanation is that mutations that alter the position and / or orientation of genes within CTDGs can be deleterious. In classic examples, like the Hox and beta-globin CTDGs, regulatory elements close to the CTDGs have been identified to be important in the expression of CTDG genes, and importantly, mutations that change the position or orientation of genes in a CTDG have phenotypic consequences (Bulger et al., 1999; Spitz, Gonzalez, Duboule, 2003).

The gibbon genome showed an interesting pattern supporting the hypothesis of CTDG conservation: if CTDGs were not conserved, the gibbon genome would have significantly lower number and length of CTDGs, given its high proportion of genomic rearrangements compared to other primates (Carbone et al., 2014). However, we found that the gibbon genome has mean fractions of clustered regions and genes across chromosomes similar to those of the rest of the primates (Fig 5). This suggests that the higher rate of genomic rearrangements in the gibbon genome did not significantly affect the length and composition of its CTDGs.

A model in which CTDG maintenance is necessary for conservation of gene regulation could explain how, because CTDGs can function as “regulatory domains” of the genes they harbor, CTDGs are maintained across evolutionary time. Similarly, large sets of co-regulated duplicated genes in close proximity would optimize their concomitant expression, because that arrangement would minimize the number of chromatin relaxation events needed for the transcription machinery to access those genes. For example, it has been shown that chromatin interactions via CTCF binding sites are necessary for the co-expression of the gene duplicates in the fibrinogen cluster (Jaimes, Fish, Neerman-Arbez, 2018).

Several additional models, not necessarily at odds with one another, could explain how CTDGs could be maintained. Right after gene duplication, it is reasonable to assume that the regulatory programs of any of the duplicates in the CTDG would activate most of the duplicates. If the increased expression of the genes in the CTDG is selected for, then a limited number of regulatory elements would be necessary. From that it follows that the regulatory elements of most duplicates in a CTDG will tend to degrade (or not being duplicated along with the gene in the first place). At that point, the CTDG would be maintained because most of the duplicates would be disregulated without the few remaining regulatory elements left. At the other end of the functional spectrum, we can imagine how CTDGs can remain duplicate-dense to increase the combinatorial potential of the CTDG. For example, the protocadherin CTDG harbors exons that can be independently regulated. In that way, many different protocadherin isoforms can be produced, increasing their functional potential as drivers of neuronal identity specification (Esumi et al., 2005; Lefebvre et al., 2012). Also, different combinations of Hox genes are expressed at different time points in the development of vertebrates, depending on the regulatory programs that are required (Holland, Garcia-Fernández, 1996). Under this model, genes in a CTDG would be maintained in close proximity to facilitate the expression of selected but different genes in the CTDG, increasing their functional repertoire.

Gene family dynamics (gene gains and gene losses) have been of great interest in evolutionary genomics research (Albalat, Cañestro, 2016; Demuth, Hahn, 2009). Assuming neutral evolution (Lynch, 2000), gene families rich in CTDGs should have a higher rate of gene duplication than gene families depleted in CTDGs because the probability of gene duplication for a gene family would be directly correlated with the number of existing gene duplicates. However, our findings suggest that duplication events have generally remained stable across gene families, regardless of the number of clustered genes in each gene family. Overall, our analysis shows that most of the gene gains per family occurred prior to the origin of mammals, which aligns with previous reports supporting that only a minority of mammal-specific homologous gene groups experienced gene duplication events in the mammalian phylogeny (Dunwell, Paps, Holland, 2017). These data, along with our experiment on the conservation of CTDGs in chromosome 19, are consistent with a model by which gene families greatly expanded before mammals diversified, and then stabilized in gene number. If this model is supported by additional data, it would call our attention to include parameters about the “genomic geography” of CTDGs in our model about the evolutionary mechanisms behind gene family evolution in particular, and evolutionary genomics in general.

## Acknowledgements

This work was conducted in part using the resources of the Advanced Computing Center for Research and Education at Vanderbilt University. This work was supported in part by the March of Dimes through the March of Dimes Prematurity Research Center Ohio Collaborative, The Burroughs Wellcome Trust Preterm Birth Initiative, the National Science Foundation (DEB-1442113 to A.R.), and the Guggenheim Foundation.

## Supplemental information

Figure S1. **GO enrichment analysis of clustered genes compared to non-clustered genes in human and mouse.** The percentage of clustered genes in each chromosome enriched for biological processes is used to color the heatmap. The GO terms represented in the heatmap belong to GO depth 6 because it is the depth in the ontology graph with the highest diversity of GO terms.

Figure S2. **Distribution of CTDGs across mammalian species.** X axis: fraction of chromosome length spanned by clusters (A), and fraction of clustered genes per chromosome (B). Y axis: density. Axes are plotted only on the human distribution to increase readability. Dotted line: fitted normal distribution, blue line: kernel density estimate, orange line: percentile 95, bottom blue ticks: data for every chromosome or scaffold, middle blue lines: data for chromosomes beyond the percentile 95. All plots are plotted on the same scale on both axes.

Figure S3. **Mean recombination rate difference between regions inside and outside of CTDGs.** The excess recombination rate inside CTDGs was calculated by subtracting the mean recombination rate outside CTDGs from the mean recombination rate inside CTDGs.

Table S1. **Genome assemblies used in the study**.

Table S2. **CTDG repertoire across selected mammalian genomes.** “numbers” sheet: summary annotation of clusters. “genes” sheet: annotation of all clustered genes and their position in each cluster

Table S3. **GO analysis.** Enriched categories at selected depth in mouse and human for biological processes

